# Vitamin D and Macrophage Polarization in Epicardial Adipose Tissue of Atherosclerotic Swine

**DOI:** 10.1101/342493

**Authors:** Palanikumar Gunasekar, Vicki J. Swier, Jonathan P. Fleegel, Chandra S. Boosani, Mohamed M. Radwan, Devendra K. Agrawal

**Author notes:** Corresponding author: (DKA).

## Abstract

Vitamin D functions as a potent immunomodulator by interacting with many immune cells however, its role in regulating inflammation in the epicardial adipose tissue (EAT) is unclear. In the EAT of atherosclerotic microswine that were fed with deficient, sufficient or supplemented levels of vitamin D, we evaluated the phenotype of the macrophages. Vitamin D treatment was continued for 12 months and serum 25(OH)D levels were measured regularly. Infiltration of M1/M2 macrophage was investigated by immunostaining for CCR7 and CD206, respectively in conjunction with a pan macrophage marker CD14. Significant difference in the number of CCR7+ cells was observed in the EAT from vitamin D-deficient swine compared to vitamin D-sufficient or -supplemented swine. Expression of CD206 correlated with high levels of serum 25(OH)D indicating a significant increase in M2 macrophages in the EAT of vitamin D-supplemented compared to -deficient swine. These findings suggest that vitamin D-deficiency exacerbates inflammation by increasing pro-inflammatory M1 macrophages, while vitamin D-supplementation attenuates the inflammatory cytokines and promotes M2 macrophages in EAT. This study demonstrates the significance of vitamin D mediated inhibition of macrophage mediated inflammation in the EAT during coronary intervention in addition to its immunomodulatory role. However, additional studies are required to identify the cellular mechanisms that transduce signals between macrophages and smooth muscle cells during restenosis in the presence and absence of vitamin D.

**Author Contribution Statement:** DKA conceived and designed the experiments; PG, JPF, MMR performed the experiments; PG, JPF, VJS analyzed and interpreted the results; PG prepared the figures and wrote the initial draft of the manuscript; CSB, DKA revised and edited the revised manuscript.

## Introduction

The epicardial adipose tissue (EAT) is the visceral fat of the heart. It lies near coronary arteries and in continuity with the myocardium. Physiologically, EAT is different from other visceral fat. EAT is a metabolically active tissue that secretes several bio-active molecules which have paracrine and vasocrine effects in the coronary artery causing vascular dysfunction and atherosclerosis (Sacks and Fain 2007; Iacobellis and Bianco 2011). EAT also secretes pro-inflammatory adipokines, including interleukin-6 (IL-6), tumor necrosis factor-alpha (TNF-α), and monocyte chemoattractant protein-1 (MCP-1) and anti-inflammatory adipokines, including adiponectin. These evidences suggest the role of EAT in establishing a chronic inflammatory environment that could lead to atherosclerosis of the adjacent coronary arteries (Mazurek et al. 2003). The tissue macrophage in EAT exists in different phenotype; pro-inflammatory M1 macrophage or anti-inflammatory M2 macrophages. Recent studies have reported that pro-inflammatory M1 macrophages are increased and anti-inflammatory M2 macrophages are decreased in EAT in patients with coronary artery disease (CAD) (Hirata et al. 2011).

Deficiency of 25-hydroxy vitamin D [25(OH)D] is found to be associated with several medical conditions, such as diabetes and insulin resistance (Scragg et al. 1995), metabolic syndrome (Hypponen et al. 2008), myocardial infarction (Scragg et al. 1990), and cardiovascular diseases (Wang et al. 2008). According to a survey conducted by the Institute of Medicine, 32% of the US population aged 1 years or above had inadequate or deficient levels of serum 25(OH)D (Looker et al. 2011). The numbers are even worse in elderly, who are the high-risk group for these diseases, with almost half of their population deficient in vitamin D (Norman et al. 2007). There are some reports implicating a detrimental effect of vitamin D-deficiency in promoting atherosclerosis (Wang et al. 2008; Norman and Powell 2005). However, most of these reports are based on epidemiological pooled data from observational studies and there is still a controversial role of vitamin D-deficiency in causation or worsening of CAD and whether vitamin D-supplements can help prevent ischemic heart diseases (Ku et al. 2013).

Vitamin D exerts its function through vitamin D receptor (VDR). Many studies have indicated a clear link between vitamin D-deficiency and cardiovascular diseases, however, the reports showed conflicting results which were attributed to the variations in the study design, bioavailability, mutations in the VDR, and the variable concentration of vitamin D in different tissues. Specific details about the benefits of vitamin D supplementation and its association with cardiovascular diseases was recently reviewed (Mozos and Marginean 2015). Gene polymorphism and genetic differences in VDR have been identified as variable factors in human population which would account for the individual differences in response to vitamin D treatment (Levin et al., 2012). The ethnic and geographical distribution of the population, and the incidence of cardiovascular diseases in these populations suggests the need for vitamin D supplementation in reducing the risk associated with different cardiovascular diseases (Levin et al., 2012). In a recent randomized placebo controlled human clinical trial, supplementation with vitamin D3 was found to effectively raise higher vitamin D levels in the blood than supplementation with vitamin D2 (Lehmann 2013). Thus, suggesting that the bioavailability of vitamin D depends on the type of isoform of vitamin D and its intake which is critical to maintaining the normal healthy levels of vitamin D in the blood.

The purpose of the present study is to identify the influence of vitamin D treatment on the EAT which reflects on the pro-inflammatory signaling in the coronary arteries that may have a profound influence on stenosis. We hypothesize that vitamin D deficiency enhances production of pro-inflammatory regulators in the EAT. Therefore, inhibition of EAT dependent regulation of pro-inflammatory signaling through vitamin D supplementation is critical to lower cellular inflammation in coronary arteries. In this study, we examined if vitamin D deficiency increases the M1/M2 macrophage ratio in EAT and thus predisposes to the development of coronary artery disease, and also evaluated whether vitamin D-supplementation would decrease M1/M2 macrophage ratio in atherosclerotic swine model.

## Materials and Method

The Institutional Animal Care and Use Committee of Creighton University approved the research protocol (IACUC#0830) and the animals were housed and maintained in the Animal Resource Facility of Creighton University, following the rules and regulations of NIH and USDA. Yucatan female microswine of 30-40 lbs were obtained from Sinclair Laboratories, Columbia, MO, USA. The microswine were fed a high cholesterol diet (Harlan Laboratories) contained the following major ingredients: 19% casein “vitamin free”, 23.5% sucrose, 23.9% corn starch, 13% maltodextrin, 4% soybean oil, 4% cholesterol, 20% chocolate mix, and 10% cellulose. Vitamin D-sufficient high cholesterol diet (Harlan, USA) was prepared with the following major ingredients: 37.2% corn (8.5% protein), 23.5% soybean meal (44% protein), 20% chocolate mix, 5% alfalfa, 4% cholesterol, 4% peanut oil, 1.5% sodium cholate, and 1% lard. The diet was either deficient in vitamin D or supplemented with 2,000 IU/d or 4,000 IU/d of Vitamin D3. Serum 25(OH)D levels were measured regularly by collecting the blood from auricular vein at baseline, before the coronary intervention and before the time of euthanasia. Serum levels of 25(OH)D were measured to determine the vitamin D status in the swine which were placed into their respective experimental groups based on the 25(OH)D levels and diets.

The animals were fed with the experimental diet for 6 months. Steps to prevent the thrombosis were taken by administering aspirin (325 mg/day) and ticlopidine (250 mg/day) to all animals three days prior to the procedure. After scrubbing and sterile draping access to the vascular system will be obtained after femoral cut down and exposure of the femoral artery. After puncturing the femoral artery, the 6F sheath was introduced to keep the artery open. After placement of sheath, an angiographic guiding catheter (6F AR2/100cm) is introduced and pushed all the way to coronary arteries on a guiding wire. Fluoroscopic evaluation is performed by administering non-ionic contrast media which is injected into the coronary arteries for fluoroscopic evaluation using C arm (OEC 9900 Elite Vas 8, GE Healthcare). Clinical “coronary-type” angioplasty catheter with a balloon size of 2.5 mm x 15 mm will be gently pushed on the guide wire, until its deflated balloon is inside the left circumflex (LCX) coronary artery, single complete balloon inflation to 10-15 atm. pressure, depending upon the vessel diameter using the inflation device with a pressure gauge (basix touchTM), then balloon is deflated, and the patency of the artery will be checked by performing angiography. Following the completion of the procedure, the catheters were removed, and the femoral artery was sutured with prolene, and the leg incision was closed with vicryl.

Angiogram and optical coherence tomography (OCT) imaging were performed using C7-XR OCT intravascular imaging system (St. Jude Medical, St. Paul, MN) at 6 months post-coronary intervention. High dose of barbiturates (Beuthanasia-D, 0.1 ml/lb, i.v.) was administered to euthanize the animal. Swine hearts were removed quickly after euthanasia and the EAT surrounding the coronary arteries were stripped and fixed in 10% formalin buffer for 24 hrs. The tissues were handled in a Sakura Tissue Tek VIP Tissue Processor and embedded in paraffin. Sections were cut at a thickness of 5µm using a microtome (Leica, Germany) and subsequently placed on slides for immunofluorescence evaluation.

## Hematoxylin and Eosin staining

Tissue sections were routinely stained with hematoxylin and eosin; Tissue sections were viewed with a Nikon™ Eclipse Ci microscope and images were photographed with a Nikon™ DS-L3 camera. The percentage of stenosis ([1-(Lumen area/Internal elastic lamina area)] * 100) of post angioplasty coronaries were grouped and analyzed statically by using one way ANOVA from Graphpad Prism6™ software.

### Immunofluorescence

Tissue sections on the slides were deparaffinized in xylene, rehydrated in ethanol, washed in double-distilled water and treated with Lab Vision™ HIER Buffer L ph9 (Thermo Fisher Scientific Inc,) at 90°C for 20 minutes. The slides were then washed three times for 3 minute each using 1x phosphate buffered saline (PBS) and then were blocked in normal horse serum from Vector™ Laboratories for 2 hours at room temperature. The primary antibodies included: mouse anti VDR (SC 13133: SCBT™), rabbit anti MCP1 (Ab 9669; Abcam™), rabbit anti TNF-α (Ab 6671; Abcam™), mouse anti CD14 (Ab 182032; Abcam™) and a combination of either rabbit anti CCR7 (Ab 32527; Abcam™) or rabbit anti CD206 (Ab 64693; Abcam™) were added overnight at 4°C. The following day, the slides were rinsed in 1x PBS and the secondary antibody (Abcam™^®^ goat polyclonal to rabbit IgG FITC Ab97199 or Invitrogen Alexa-Fluor^®^ 594 goat anti-mouse IgG) was added and incubated on the slides for 2 hours at room temperature. Sections were washed in 1x PBS and nuclei were counterstained with Vector™ laboratories DAPI. Pictures were taken with Olympus BX-51 epifluorescent microscope and images were photographed with an Olympus DP71 camera. The images were then quantified using CellSens™ Dimension software. A region of interest (ROI) was drawn around individual nuclei. The software quantified the protein expression in mean intensity value for each nucleus. The 40x images were quantified from each EAT section per swine. The data from each experimental group of swine (vitamin D-deficient, vitamin D-sufficient, and vitamin D-supplemented) were grouped and analyzed statistically by using one-way ANOVA using GraphPad Prism6™ software. For Double immunofluorescence staining the 40x images were manually analyzed by two observers and the average of the positive cell count was used for statistical analysis. There was less than 5% variation between the observers.

### Statistical analysis

The immunofluorescence data were analyzed by GraphPad Prism 6.0 (GraphPad Software, Inc). The values are presented as mean ± SEM. One-way ANOVA with Turkey’s multiple comparison test was used to analyze significant differences in mean fluorescence index (MFI) in each cytokine within the experimental groups, and positive cell count for CCR7^+^ and CD206^+^ cells in the experimental groups. The data in Fig 2I, 6A, and 6B were analyzed via non-linear regression and fit with exponential decay lines. A p value of <0.05 was considered statistically significant.

## Results

### Evaluation of serum 25(OH)D levels in different animal groups

Animals were classified into three groups based on the measured serum 25(OH)D levels. Group 1. Animals with ≤20 ng/ml of 25(OH)D were grouped as vitamin D-deficient, Group 2. Animals with the serum levels of 25(OH)D ranging between 30-44 ng/ml were grouped as vitamin D sufficient, and Group 3. Animals with more than 44 ng/ml of 25(OH)D in serum were grouped as vitamin D-supplemented. Each of these experimental group animals had a sample size of 3 for a total of 9 swine in this study. The mean serum 25(OH)D levels observed in the three experimental groups were: deficiency 17.1 ± 1.6 ng/ml, sufficient 37.4 ± 3.7 ng/ml, and supplemented 58.06 ± 4.4 ng/ ml. Fig 1, shows the mean serum 25(OH)D levels in different animal groups with SEM. Serum 25(OH) D level in vitamin D-deficient was lower compared to vitamin D-sufficient (p< 0.014) and vitamin D-supplemented swine (p< 0.0004). The serum 25(OH) D level in vitamin D-sufficient was lower compared to vitamin D-supplemented swine (p<0.013).

**Fig 1:**
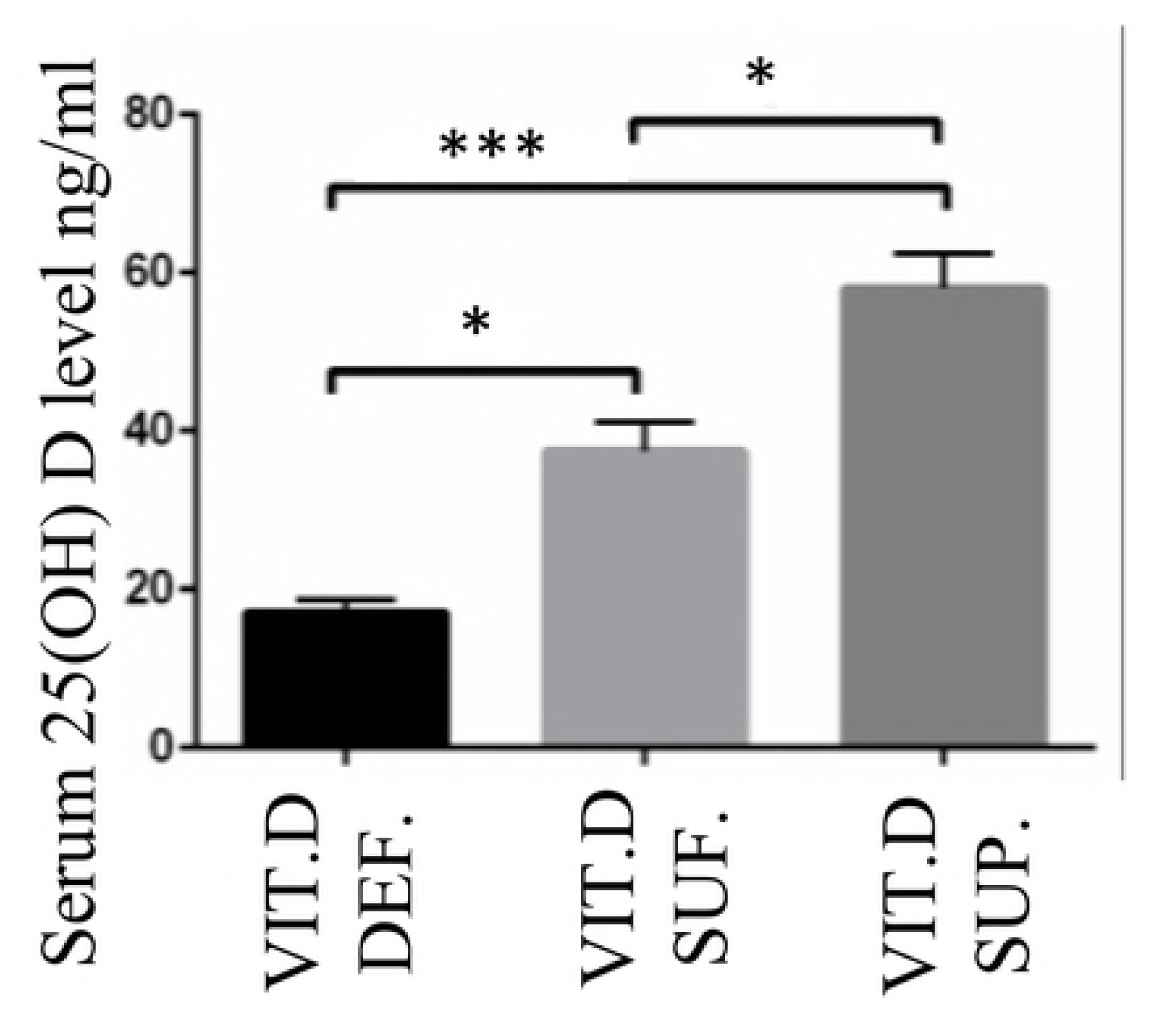
Evaluation of serum 25(OH)D levels in different animal groups. Serum 25(OH) D level in vitamin D-deficient (Vit.D.Def.), vitamin D-sufficient (Vit.D.Suf.) (p< 0.014) and vitamin D-supplemented swine (Vit.D.Sup.) (p< 0.0004).

### Hematoxylin and eosin stain

The area of neo-intimal region (Fig 2A-C) following angioplasty of coronary arteries was greater in vitamin D-deficient compared to vitamin D-sufficient and vitamin D-supplemented swine. The percentage restenosis (Fig 2G) in post angioplasty coronary arteries was greater in vitamin D-deficient swine (62.41 ± 4.067%) compared to vitamin D-sufficient (37.90 ± 4.264%) (p<0.003) and to vitamin D-supplemented swine (31.94 ± 2.780%) (p<0.0005).

**Fig 2:**
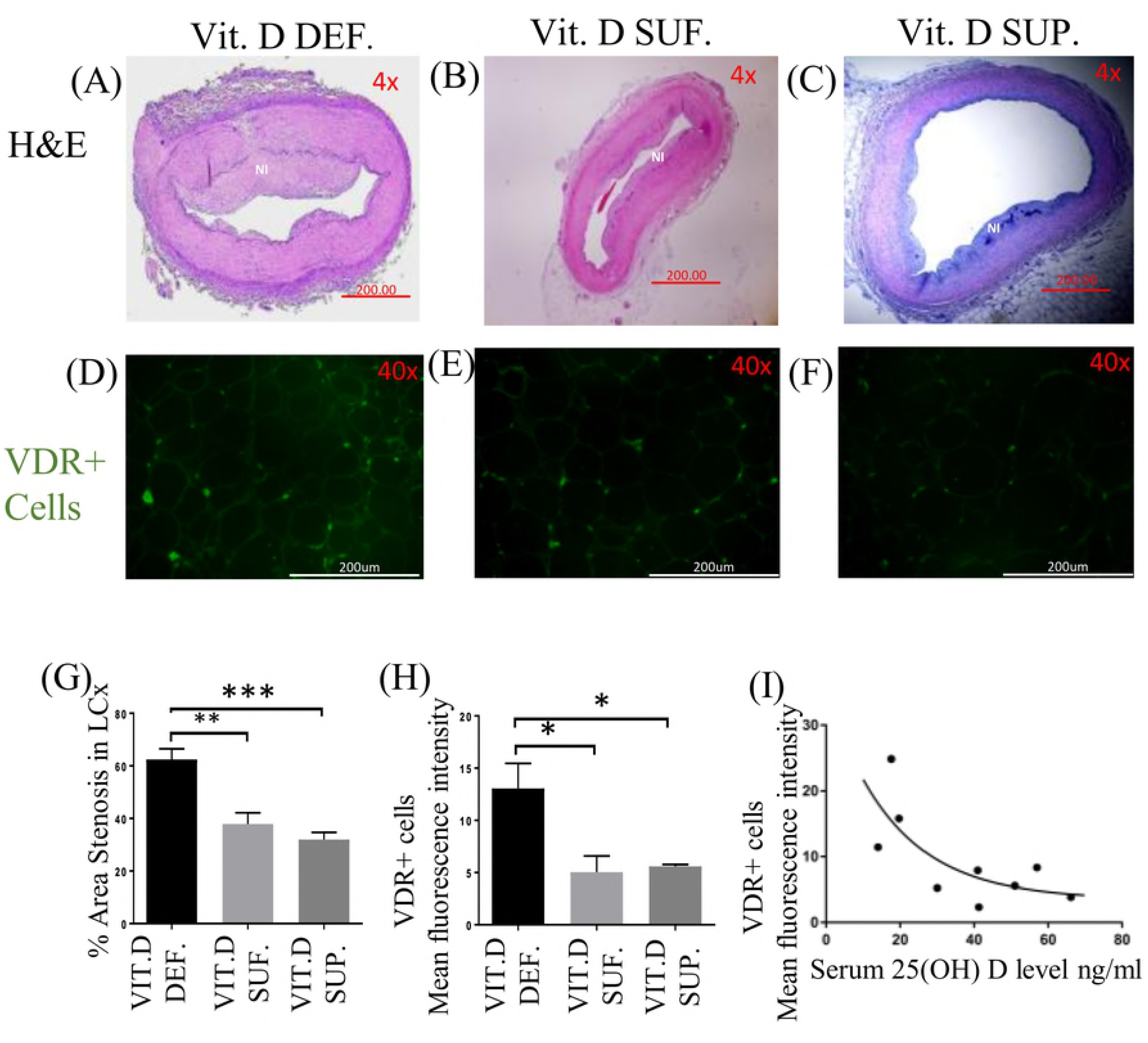
Hematoxylin and Eosin (H&E) staining and expression of vitamin D receptor (VDR). (A-C) shows the H&E staining of post-balloon angioplasty site of the left circumflex coronary artery (LCx); (D-F) shows the effect of vitamin D on vitamin D receptor (VDR) expression in EAT of: vitamin D-deficient (Vit.D.Def.), vitamin D-sufficient (Vit.D.Suf.) and vitamin D-supplemented (Vit.D.Sup.) groups; (G) shows the percent stenosis. (n=3 in each treatment group); ***p<0.0005, **p<0.003. (H) The mean fluorescent intensity (MFI) was measured for VDR+cells in each treatment group. (n=3 in each experimental group); *p<0.04. (I) A scatter plot showing the MFI of VDR with change in the serum 25(OH) D level.

### Expression of VDR in EAT

VDR expression in EAT (Fig 2D-F) was significantly higher in the vitamin D-deficient (MFI 13.06 ± 2.40) swine compared to vitamin D-sufficient (MFI 5.04 ± 1.54) (p=0.03) and vitamin D-supplemented (MFI 5.59 ± 0.17) (p=0.04) swine (Fig 2H). But, no significantly difference was seen between vitamin D-sufficient (MFI 5.04 ± 1.54) and vitamin D-supplemented (MFI 5.59 ± 0.17) (p=0.9) swine. Serum 25(OH)D levels was inversely related to VDR expression in EAT. Fig 2I illustrates the relationship between VDR expression in the EAT, measured by fluorescent intensity, and serum 25(OH)D levels in each pig. A non-linear relationship is described by the exponential decay line as shown. The R-squared value is 0.5328.

### **Expression of MCP1 and TNF**-**a in EAT**

The TNF-α expression (Fig 3A-C) was significantly greater in EAT of vitamin D-deficient (MFI 107.0 ± 2.34) swine compared to vitamin D-sufficient (MFI 60.15 ± 2.15) (p<0.0001) and vitamin D-supplemented (MFI 48.35 ± 1.66) (p<0.0001) swine. There was also significantly increase in the expression of TNF-α in the EAT of sufficient (MFI 60.15 ± 2.15) swine compared to a supplemented swine (MFI 48.35 ± 1.66) (p=0.001) (Fig 3G).

**Fig 3:**
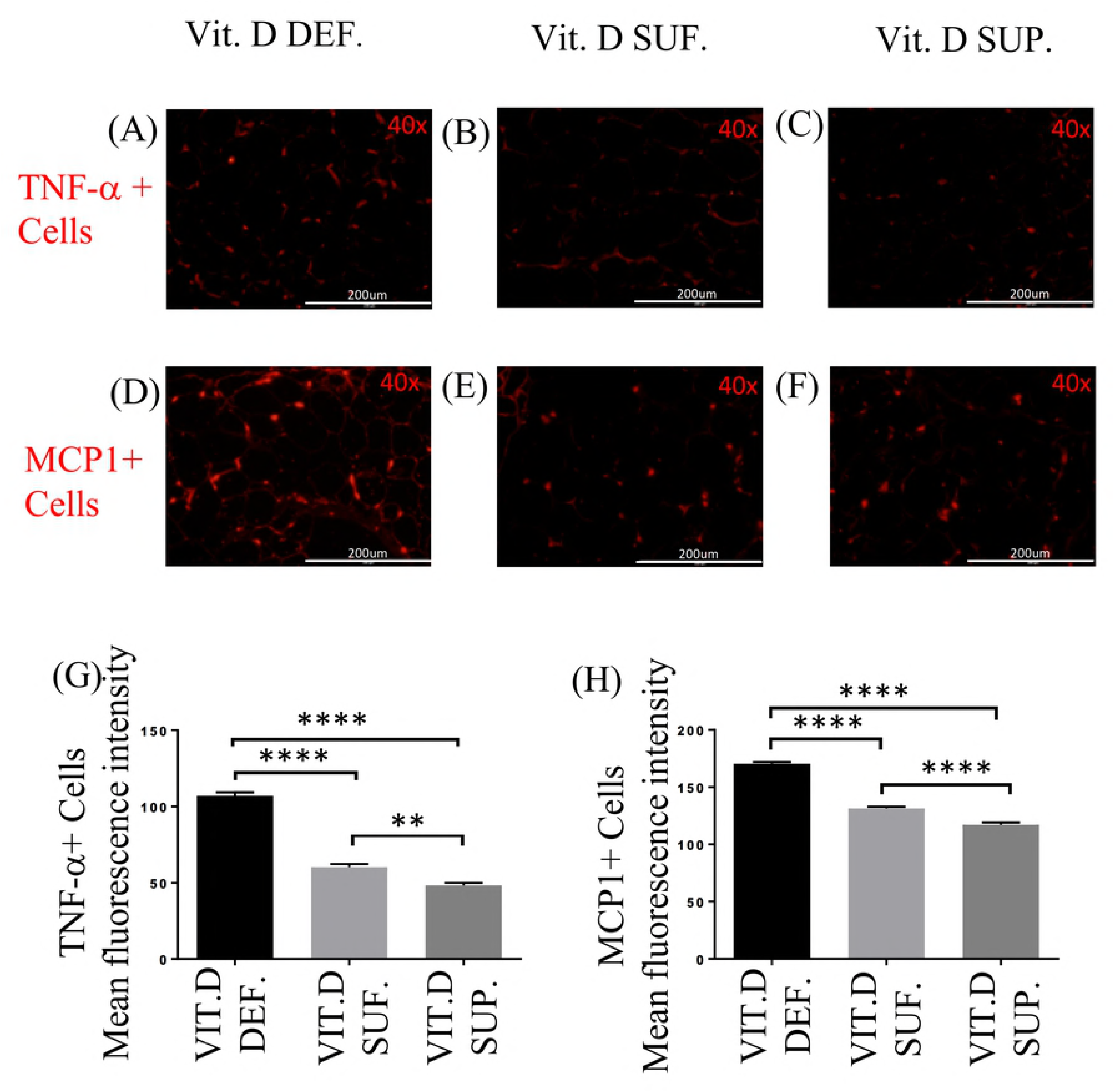
Effect of vitamin D on TNF-α and MCP1 expression. Comparison of Immunofluorescence staining for TNF-α (A-C) and MCP1 (D-F) cells in the EAT of: vitamin D-deficient (Vit.D.Def.), vitamin D-sufficient (Vit.D.Suf.) and vitamin D-supplemented (Vit.D.Sup.) groups. (G) The mean fluorescent intensity (MFI) values for TNF-α^+^cells were measured for each treatment group. (H) The MFI values for MCP1 cells were measured for each treatment group. Data show the MFI ± SEM (n=3 in each treatment group); **p <0.0012. ****p <0.0001.

The MCP1 expression in EAT (Fig 3D-F) was significantly greater in vitamin D-deficient (MFI 170.4 ± 1.68) swine compared to vitamin D-sufficient (MFI 131.4 ± 1.51) (p<0.0001) and vitamin D-supplemented swine (MFI 117.1 ±2.04) (p< 0.0001). There was also significantly difference in MFI for MCP1^+^ cells in the EAT between vitamin D-sufficient (MFI 131.4 ± 1.51) and vitamin D-supplemented swine (MFI 117.1 ±2.04) (p< 0.0001) (Fig 3H).

### CD14/CCR7 and CD14/CD206 expression in EAT

The double immunostaining for CD14 and CCR7 (Fig 4A-I) was performed to identify M1 macrophages in the EAT. There were significantly more number of cells expressing CCR7 (M1 macrophages) in the EAT of vitamin D-deficient swine (9.86 ± 0.32) compared to vitamin D-sufficient swine (3.20 ± 0.24) (p<0.0001); and to vitamin D-supplemented swine (2.43 ± 0.19) (p<0.0001). The cell counts of CCR7^+^ cells in the EAT of vitamin D-supplemented swine were not statistically different from those in vitamin D-sufficient swine (p =0.11) (Fig 4J).

**Fig 4:**
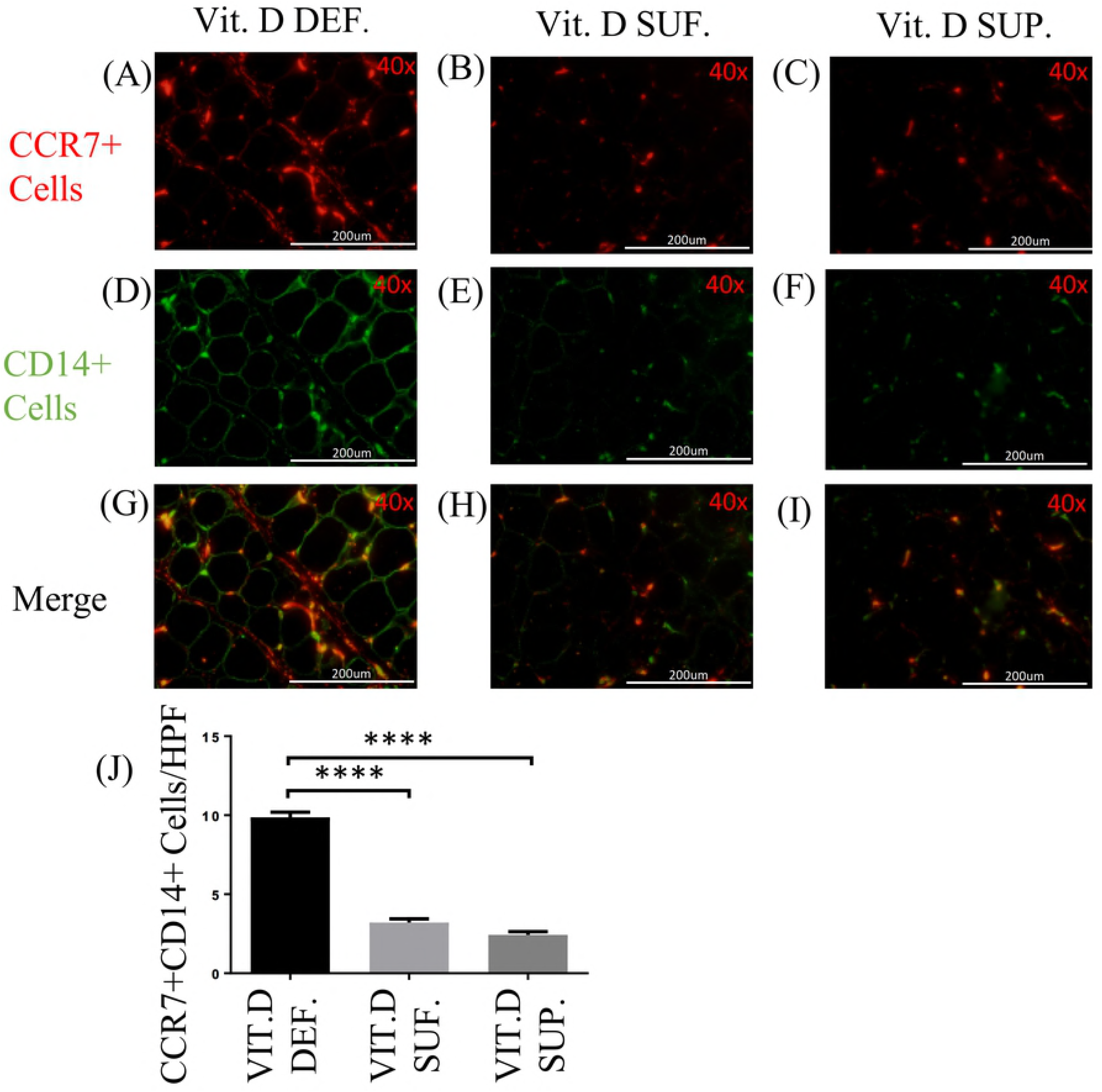
Effect of vitamin D on CD14/CCR7 expression. (A-I) Double immunofluorescence staining for CCR7 and CD14 in EAT of: vitamin D-deficient (Vit.D.Def.), vitamin D-sufficient (Vit.D.Suf.) and vitamin D-supplemented (Vit.D.Sup.) groups. (A-C) Cy-3 (red) images are CCR7^+^ Cells. (D-F) FITC (green) images are all CD14^+^ Cells. (J) Data reveal that more cells were expressing CCR7 in vitamin D-deficient swine and lower in vitamin D-supplemented swine than with vitamin D-sufficient swine. (n=3 in each treatment group); ****p <0.0001.

The double immunostaining with CD14 and CD206 (Fig 5A-I) was performed to identify macrophages of the M2 phenotype in the EAT. The density of CD206^+^ cells in the EAT of vitamin D-deficient swine (mean 4.86 ± 0.41) was found to be significantly less than those in vitamin D-sufficient swine (mean 10.63 ± 0.55) (p<0.0001) and in vitamin D-supplemental swine (mean 12.60 ± 0.66) (p<0.0001). The CD206+ M2 macrophages in the EAT were significantly greater in the vitamin D-supplemented than vitamin D-sufficient swine (p<0.05) (Fig 5J).

**Fig 5:**
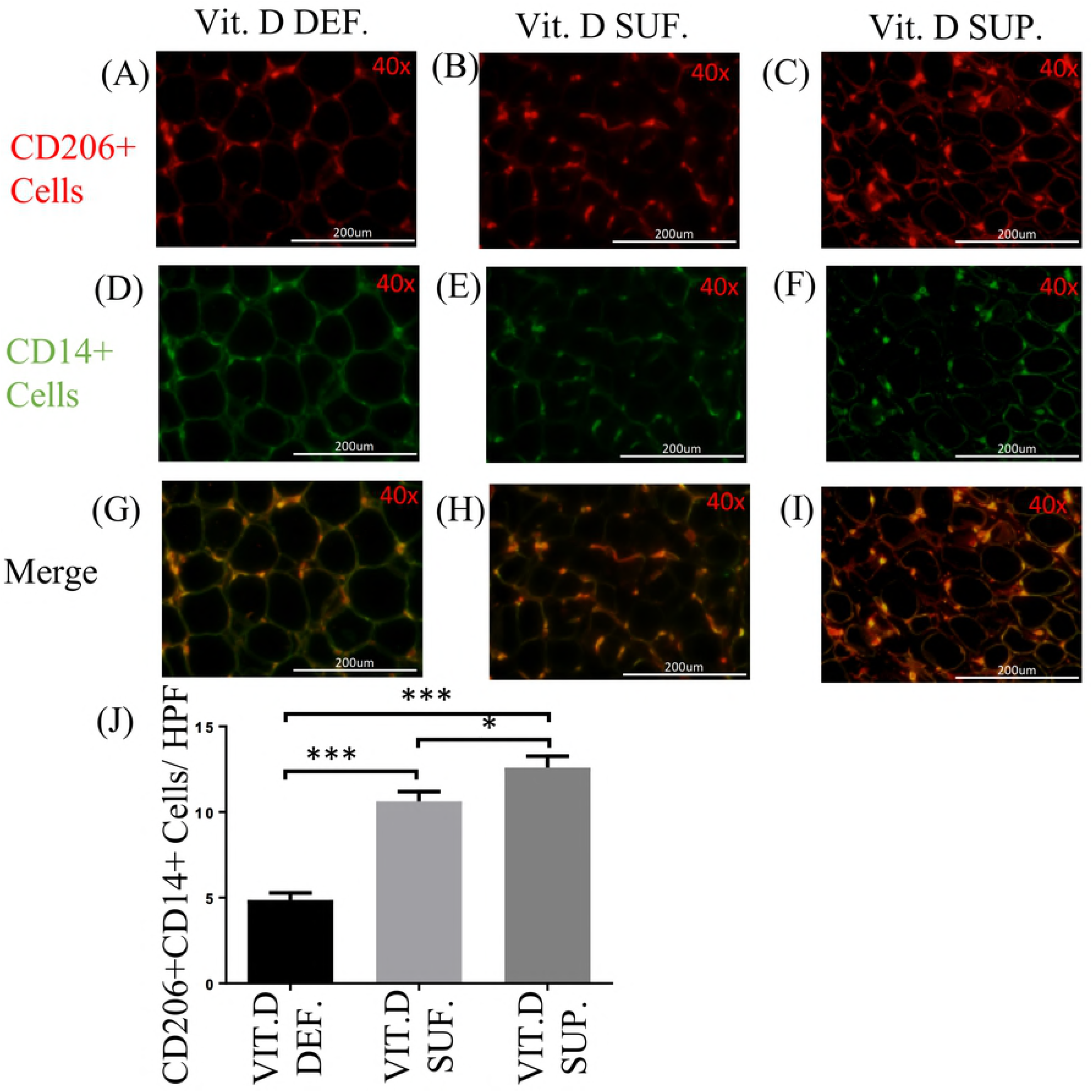
Effect of vitamin D on CD14/CD206 expression. (A-I) Double immunofluorescence staining for CD206 and CD14 in EAT of: vitamin D-deficient (Vit.D.Def.), vitamin D-sufficient (Vit.D.Suf.) and vitamin D-supplemented (Vit.D.Sup.) groups. (A-C) Cy-3 (red) images are CD206^+^ Cells. (D-F) FITC (green) images are all CD14^+^ Cells. (J) Data reveals that few cells were expressing CD206 in vitamin D-deficient swine and more in vitamin D-supplemented swine than with vitamin D-sufficient swine. (n=3 in each treatment group); *p <0.05, ****p <0.0001.

We compared the relationship between serum 25(OH)D levels with inflammatory parameters in the EAT of all swine. The presence of M1 and M2 macrophages in the EAT of each animal is shown in Fig 6A. Decrease in M1 macrophages with a concomitant increase in M2 macrophages was observed with the increase in the serum 25(OH)D levels. The level of serum 25(OH)D at which the net polarization state of the macrophages switched from a predominately M1 state to a predominately M2 state occurred around 30 ng/mL of serum 25(OH)D. After this point, the rate of macrophage phenotypic change stabilized. This data was best modeled by exponential decay in which the R-squared value for the M1 macrophage polarization state is 0.83 and the M2 macrophage polarization state is 0.79. Next, in Fig 6B we modeled the relationship between M1 and M2 macrophage polarization by plotting the ratio of the number of M1 macrophages in the EAT to the number of M2 macrophages in the EAT of the same animal. Once again, the data was represented by an exponential decay relationship with the R-squared value of 0.90.

**Fig 6:**
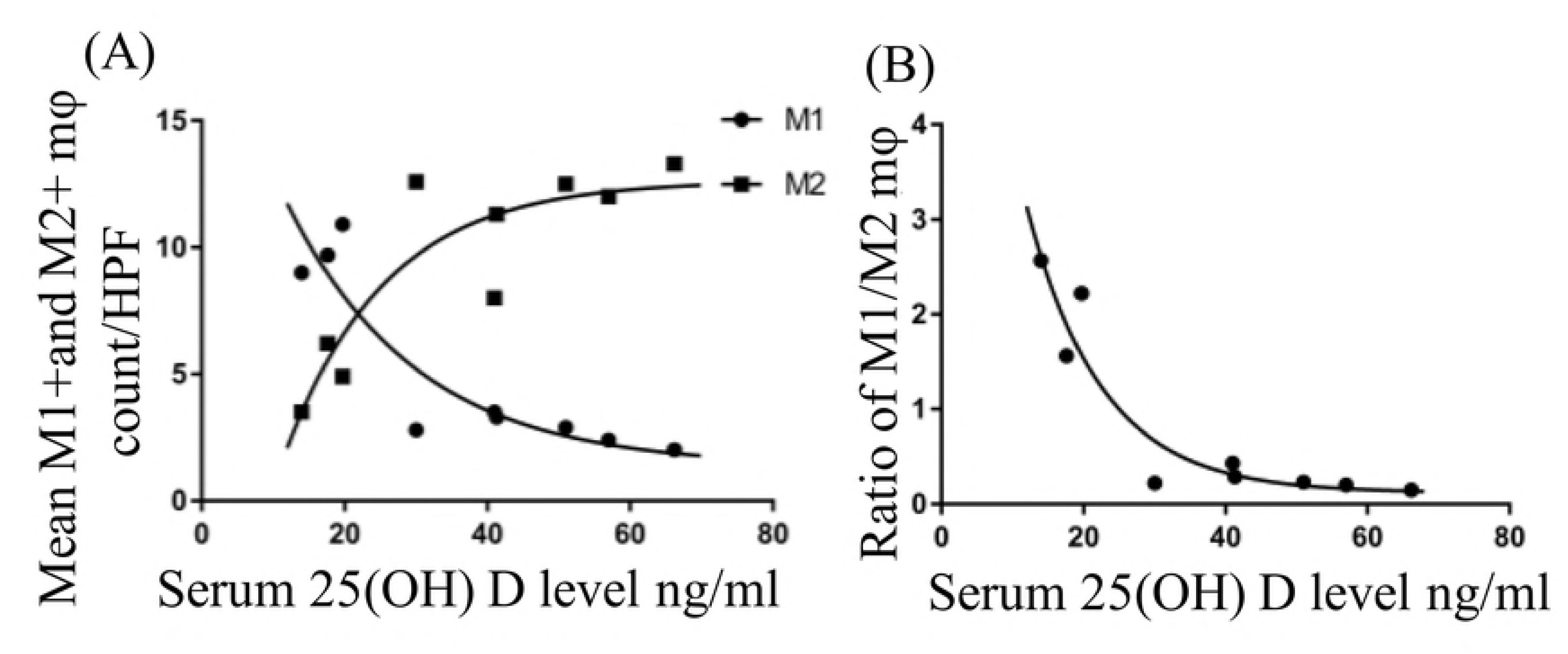
A scatter plot of the M1 and M2 macrophage. (A) A representation of the M1 and M2 macrophage count in the epicardial adipose tissue (EAT) of each animal against their serum 25 (OH)D levels. The solid circle (●) represents M1 macrophage count for the swine under 10 high power field (HPF). The solid square (■) represents M2 macrophage count for the swine under 10 HPF. (B) A scatter plot of the M1/M2 macrophage ratio against their serum 25 (OH)D level in each animal.

With increase in the serum 25(OH)D levels, the expression levels of MCP1, TNF-α, CCR7 and CD14 in the EAT were gradually decreased. These results suggest a proactive role of vitamin D as an immunomodulator in the EAT.

## Discussion

Vitamin D is a critical mediator in calcium-phosphorus homeostasis in bone metabolism as well as it is a potent immunomodulator and decreases the production of pro-inflammatory cytokines, while increasing anti-inflammatory mediators. The cytokine milieu determines the recruitment of either M1 (pro-inflammatory) or M2 (anti-inflammatory) macrophages (Myszka and Klinger 2014). There are studies demonstrating significant effect of vitamin D deficiency in the pathogenesis and exacerbation of CAD (Dozio et al. 2015; Verdoia et al. 2014; Seker et al. 2014). Also, vitamin D deficiency associates with increased inflammatory markers in EAT and was shown to induce hypertrophy in cardiac myocytes (Gupta et al. 2012).

Vitamin D acts through VDR, which is a member of steroid hormone nuclear receptor superfamily. This regulates transcription of many target genes. VDR is expressed in most of human tissues, including macrophages, pancreatic beta-cells, epithelial cells, osteoblast. The presence of VDR in most of the tissues provides a mechanistic link between vitamin D deficiency and the pathophysiology of the disorders. Recently, increased VDR expression in the medial smooth muscle cells of the coronary artery was observed in vitamin D deficient rats (Hadjadj L et al., 2018). Although the exact mechanism is still unclear, the report suggests that the insulin-mediated coronary arteriole relaxation was affected in the absence of vitamin D and this correlates with increased expression of VDR.

In this study, we demonstrated the immunomodulatory effect of vitamin D supplementation in EAT of hypercholesterolemic swine. The findings of the M1:M2 ratio in our study demonstrate that there was a significant increase in the number of M1 macrophages in vitamin D-deficient swine. Vitamin D sufficiency or supplementation significantly increases the density of M2 macrophages in EAT. Activated M2 macrophages show increased expression of CD206 (mannose receptor) and are involved in tissue repair (Folias et al. 2014). Mannose receptor (CD206) expression is upregulated by IL-10, thereby recruiting more M2 macrophages and reduce the inflammatory process (Ding et al. 2012). The M1 macrophages are characterized by the cell surface expression of CCR7 that plays a role in the activation of inflammatory M1 macrophages and migration of T lymphocytes from blood to inflamed tissues. Signals mediated through CCR7 result in activation and polarization of T cells to Th1 cells (Noor and Wilson 2012). In a study of T cells in mice, a notable inhibitory effect of vitamin D was reported on Th1 cells with a concomitant increase in Th2 cell development (Boonstra et al. 2001).

In this present study, we found that swine with serum 25(OH)D levels above 30 ng/ml had a decrease in inflammatory M1 macrophages and an increase in anti-inflammatory M2 macrophages in the EAT. The difference in macrophage phenotype could be mediated through the inhibition of IFN-γ by vitamin D and the inhibition of M1 macrophage differentiation by IL-10, which is secreted by M2 macrophages. The inhibition of IFN-γ subsequently reduces macrophage differentiation of the M1 type. Also, as the ratio decreases, the number of M2 macrophages increase. M2 macrophages produce IL-10 which inhibits the differentiation of M1 macrophages. This is supported by our study showing a positive relationship between serum 25(OH)D levels and M2 macrophages, and by the data that M1 macrophages were decreased with increasing levels of serum 25(OH)D levels. These findings suggest that vitamin D has an impact by macrophage polarization towards M2 phenotype. The major source of IL-10 is from M2 macrophages present in the EAT (Norman and Powell 2014; Ouyang et al. 2011). This could be further supported by the comparison of the ratio of M1:M2 in each treatment group.

EAT acts like an endocrine organ because it is the source of several bioactive proteins. For example, proatherogenic (IL-6, TNF-α, MCP1) and anti-inflammatory (IL-10, adiponectin) adipokines originate in the EAT (Iacobellis 2015). Dozio et al. reported increased gene expression of inflammatory molecules in the EAT of vitamin D-deficient patients with CAD compared to vitamin D-sufficient and -insufficient patients with CAD (Dozio et al. 2015). In our study, the levels of TNF-α and MCP1 decreased significantly from vitamin D-deficient to vitamin D-sufficient but not to the same extent from vitamin D-sufficient to vitamin D-supplemented groups. This can be explained by the ratio of M1:M2 that decreased dramatically from vitamin D-deficiency to vitamin D-sufficiency, but the extent at which it decreases from vitamin D sufficient to vitamin D supplemented group was less. This indicates that the TNF-α and MCP1 levels in EAT are influenced by the M1:M2 ratio. Our results are in accordance with the report by Giulietti et al. (Giulietti et al. 2007) where the immunomodulatory role of vitamin D3 was executed through the down-regulation of TNF-α in monocytes of type 2 diabetic patients. Vitamin D has been shown to inhibit the production of IL-6, IL-8 and IFN-γ by peripheral blood mononuclear cells in psoriatic patients (Inoue et al. 1998) and also decreases production of pro-inflammatory cytokines, IL-6 and TNF-α, by inactivating p38 kinase in human monocytes (Zhang et al. 2012).

Fat tissues are the major storage site for vitamin D in the body (Lawson et al. 1986; Mawer et al. 1972). In this study, we did not measure the vitamin D concentration in EAT, as there was a strong association between EAT and serum vitamin D levels based on a study by Blum et al. who demonstrated that serum and subcutaneous fat tissue vitamin D3 concentrations are positively correlated (Blum et al. 2008). Didriksen and colleagues demonstrated that subcutaneous adipose tissue could store large amount of vitamin D upon supplementation (Didriksen et al. 2015). It is not known about how the storage of vitamin D in adipose tissue is regulated and whether the stored vitamin D plays an active role.

## Conclusion

In conclusion, vitamin D deficiency increases the pro-inflammatory adipokine expression and the recruitment of M1 macrophages in the EAT. Vitamin D supplementation decreases inflammatory processes in the EAT and promotes the anti-inflammatory responses in the EAT. Fig 7 summarizes the key findings and interpretations from our study. Considering the epidemiological data showing the protective role of vitamin D in several inflammatory diseases, this study further supports the beneficial effect of vitamin D supplementation in reducing the burden of coronary artery diseases. To our knowledge this is the first report that shows vitamin D3 supplementation would regulate inflammatory responses in the EAT and alter macrophage polarization.

**Fig 7:**
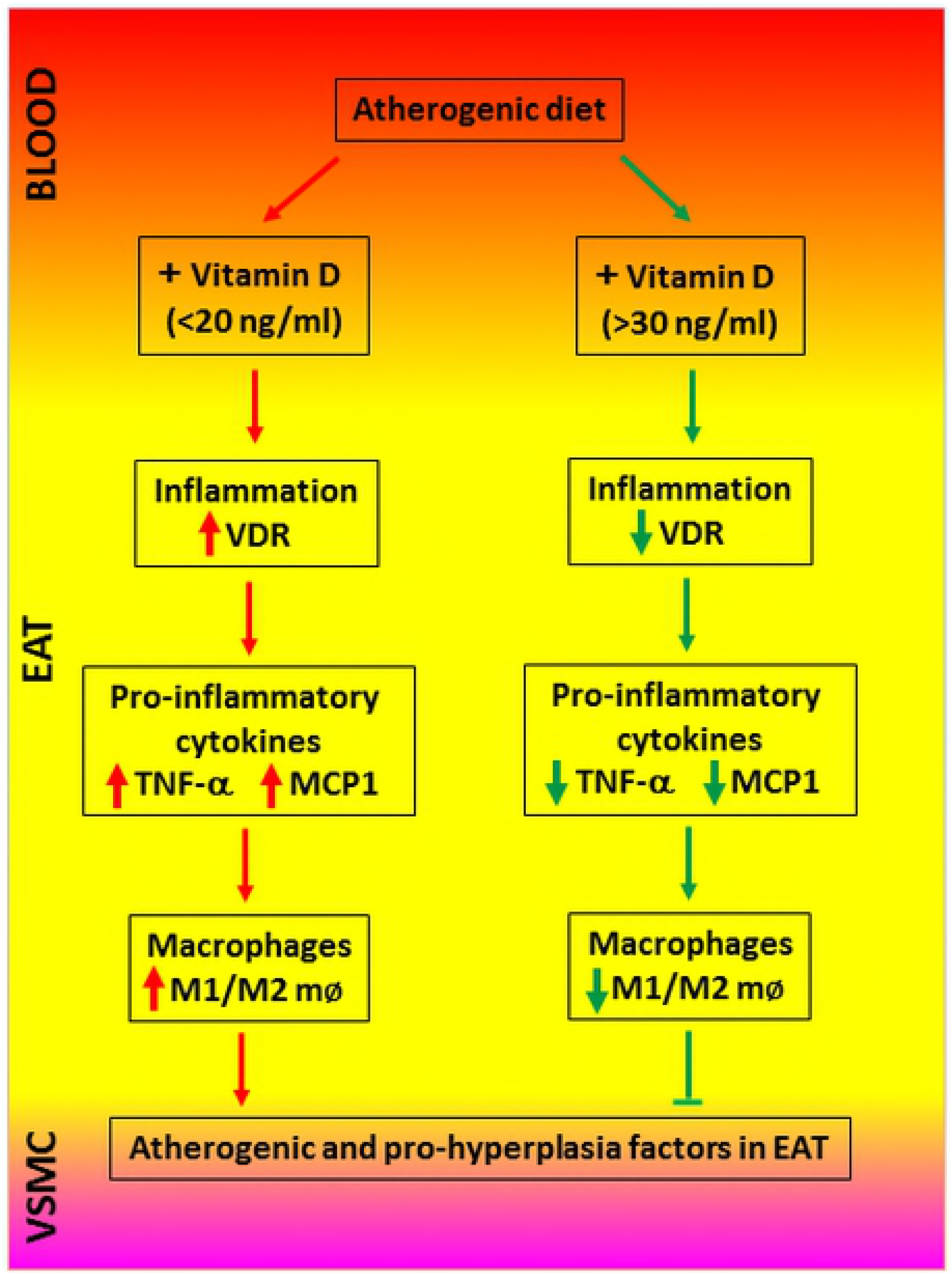
Pictorial representation of the effects of vitamin D in regulating pro-inflammatory signaling in the epicardial adipose tissue (EAT). Vascular smooth muscle cells (VSMC).

## Acknowledgments

The authors thank their colleagues in the laboratory in handling and monitoring swine at various stages of the project.

